# scAnnotate: an automated cell type annotation tool for single-cell RNA-sequencing data

**DOI:** 10.1101/2022.02.19.481159

**Authors:** Xiangling Ji, Danielle Tsao, Kailun Bai, Min Tsao, Li Xing, Xuekui Zhang

## Abstract

**Motivation:** Single-cell RNA-sequencing (scRNA-seq) technology enables researchers to investigate a genome at the cellular level with unprecedented resolution. An organism consists of a heterogeneous collection of cell types, each of which plays a distinct role in various biological processes. Hence, the first step of scRNA-seq data analysis is often to distinguish cell types so they can be investigated separately. Researchers have recently developed several automated cell type annotation tools, requiring neither biological knowledge nor subjective human decisions. Dropout is a crucial characteristic of scRNA-seq data widely used in differential expression analysis. However, dropout information is not explicitly used by any current cell annotation method. Fully utilizing dropout information for cell type annotation motivated this work.

**Results:** We present scAnnotate, a cell annotation tool that fully utilizes dropout information. We model every gene’s marginal distribution using a mixture model, which describes both the dropout proportion and the distribution of the non-dropout expression levels. Then, using an ensemble machine learning approach, we combine the mixture models of all genes into a single model for cell-type annotation. This combining approach can avoid estimating numerous parameters in the high-dimensional joint distribution of all genes. Using fourteen real scRNA-seq datasets, we demonstrate that scAnnotate is competitive against nine existing annotation methods. Furthermore, because of its distinct modelling strategy, scAnnotate’s misclassified cells are very different from competitor methods. This suggests using scAnnotate together with other methods could further improve annotation accuracy.

**Availability:** We implemented scAnnotate as an R package and made it publicly available from CRAN.

**Contact:** Xuekui Zhang: and Li Xing

## 1 Introduction

Every biological process in the human body relies on the coaction of numerous cell types, each with its own designated function. Cell identification is thus crucial in studying biological phenomenons and developing medical practices; pathology, for example, hinges on the accuracy of this task. Although standard immunophenotyping methods are widely practiced for cell identification, their heavy reliance on the manual selection of antibodies, markers and fluorochromes renders new and rare cell types particularly difficult to identify (Gown, 2008; Leach *et al*., 2013). Conversely, newly developed single-cell RNA sequencing (scRNA-seq) technologies (Tang *et al*., 2009) have heightened the detail with which we can examine cell composition by offering an unprecedented resolution of gene expression at the cellular level. The recent surge of available scRNA-seq data allows for increased accuracy in several aspects of genomic data analysis, including cell annotation (Chen *et al*., 2019; Diaz-Mejia *et al*., 2019; Artegiani *et al*., 2017).

Cell-type annotation using scRNA-seq data enables researchers to distinguish various types of cells from heterozygous populations, then investigate each cell type separately and learn their interactions. Hence, cell-type annotation is often the first s tep o f s cRNA-seq d ata analysis, which has led to a recent surge of methods developed for this task. Pasquini *et al*. Pasquini *et al*. (2021) discuss 24 scRNA-seq cell-type annotation methods developed in the last five years. The most popular cell-type annotation approach was clustering analysis followed by manual annotation. The most important advantage of such an approach is that it does not require training a model using another ‘annotated’ scRNA-seq dataset. However, such unsupervised machine learning approaches have a critical issue; namely, they require users to manually label the cell types for each cluster of cells. The manual decisions need special biological knowledge and are subjective to researchers’ individual opinions, which can be time-consuming and inconsistent. In the last few years, a huge amount of scRNA-seq data was generated and made publicly available. These rich resources made it easier and easier to identify suitable data for training supervised machine learning models to annotate new scRNA-seq data. Recently, many supervised machine learning methods have been developed for cell-type annotation.

Currently, the discriminative classification approach dominates supervised machine learning methods for cell-type annotation. This discriminative classification approach models the distribution of cell types conditional on genomic data. For example, CaSTLe (Lieberman *et al*., 2018) employs an XGBoost (Chen and Guestrin, 2016) classification model and SingleCellNet (Tan and Cahan, 2019) trains a Random Forest classifier on discriminating gene pairs. CHETAH (de Kanter *et al*., 2019) and scClassify (Lin *et al*., 2020) construct hierarchical classification trees and evaluate the correlation of query cells to reference cell types or apply an ensemble of weighted kNN classifiers, respectively. In SingleR (Aran *et al*., 2019), the Spearman rank correlations of query cells to reference samples are used in an altered kNN classification algorithm. Similarly, scmap (Kiselev *et al*., 2018) classifies cells by measuring their similarity to either the centroids of reference clusters (scmap-cluster) or by kNN cell annotation (scmap-cell). Finally, scPred (Alquicira-Hernandez *et al*., 2019) reduces the reference data’s dimensionality using PCA and applies a Support Vector Machine model for classification. The common unwanted characteristic of discriminative classification methods is that they do not utilize the distribution of genomic data. However, the distribution of genomic data carries key features of scRNA-seq data, which should be helpful for cell-type annotation. For example, “dropout” is the well-known sparsity issue characterized by the excessive amount of zero counts in scRNA-seq data, arising from technical limitations in detecting moderate or low gene-expression levels in cells of the same type (Hicks *et al*., 2018). Various imputation methods have been developed to remove dropouts from data, such as SAVER (Huang *et al*., 2018), scImpute (Li and Li, 2018), and DrImpute (Gong *et al*., 2018). However, imputation could generate false positive signals within the data due to the intrinsic circularity of current scRNA-seq expression recovery practices (Andrews and Hemberg, 2018). Furthermore, the proportion of dropouts can provide helpful information for cell annotation. We, therefore, prefer to utilize this information instead of removing it.

To fully utilize the unique characteristics of scRNA-seq genomic data, we investigate the generative classification approach which models the distribution of genomic data conditional on cell type. Such an approach focuses on distributions of genomic data in different cell types, and annotates cells using the Bayesian theorem. To the best of our knowledge, scID (Boufea *et al*., 2020) is the only cell-type annotation method based on a generative classifier. scID uses Fisher’s linear discriminant analysis (LDA) to distinguish the characteristic genes of pre-determined cell clusters. LDA assumes that genomic data follow a multivariate normal distribution, which might over-simplify the complexity of the data. Furthermore, from data with limited sample sizes, it is hard to precisely estimate numerous parameters in the high-dimensional covariance matrix of the assumed multivariate normal distribution.

In this paper, we propose a novel generative classifier for automated cell-type annotation, scAnnotate. We focus on addressing the two critical challenges of scRNA-seq data as discussed above: the curse of high dimensionality (as discussed in LDA) and explicitly modelling dropout. To address the curse of high dimensionality, we use every gene to make a classifier and consider it as a ‘weak’ learner, and then use a combiner function to ensemble ‘weak’ learners built from all genes into a single ‘strong’ learner for making the final decision. To select a gene’s distribution that explicitly models the excessive zero counts in each weak learner, we borrow the idea from differential expression (DE) analysis of scRNA-seq data. The literature of DE analysis is well-established, with many methods that focus on modelling excessive zero counts. For example, Kharchenko *et al*. Kharchenko *et al*. (2014) introduced a Bayesian approach to scRNA-seq DE analysis in which non-zero counts are modelled using a Negative Binomial distribution, and zero counts are modelled with a low-magnitude Poisson process. DEsingle Miao *et al*. (2018) is another scRNA-seq DE analysis tool that uses the Zero-Inflated Negative Binomial (ZINB) distribution. However, after batch effect removal and other preprocessing, scRNA-seq data are often no longer integers and hence are not suitable for the ZINB model. Furthermore, recent benchmark studies did not show any clear advantage of the ZINB model in DE analysis of scRNA-seq data (Soneson and Robinson, 2018). We, therefore, model gene expression levels as a continuous variable. MAST (Finak *et al*., 2015) joint models the proportion of dropouts and the distribution of non-dropouts using a hurdle regression model, which is one of the most popular DE analysis softwares, and has shown great performance in benchmark studies (Soneson and Robinson, 2018). Inspired by MAST, we joint model the proportion of dropouts and the gene expressions of non-dropouts by a two-component mixture model. We tried various distributions to model the non-dropout component in the mixture model and found empirically that the lognormal distribution works best for most of the data that we explored. In the Discussion, we also discuss two alternative distributions implemented in our software that are useful in particular situations. In the rest of this paper, we will introduce the details of the scAnnotate method and use real scRNA-seq datasets to compare its classification performance with nine other scRNA-seq annotation methods based on supervised machine learning algorithms.

## 2 Materials and methods

We introduce scAnnotate, an automated cell type annotation tool. scAnnotate is entirely data-driven, meaning that it requires training data to learn the classifier but does not require biological knowledge or subjective decisions from the user. It consists of three steps: preprocessing training and test data, model fitting on training data, and cell classification on test data. The classification model in the last step uses an ensemble machine learning approach involving many weak learners and a combiner function to integrate the outputs of the weak learners into a single strong learner. Each weak learner is a classifier based on a mixture model for the expression level of one gene. The combiner is a weighted average of all weak learners’ outputs. The weights can be either learned from training data with at most one rare cell population, or pre-specified as equal weights on the training data with at least two rare cell populations. In this study, we defined a rare cell population as a cell population with less than 100 cells in a given dataset. scAnnotate handles data with varying numbers of rare cell populations differently. An illustration of its workflow is shown in Figure 1 (dataset with at most one rare cell population) and Figure 2 (dataset with at least two rare cell populations). Details of each element of scAnnotate will be discussed in the rest of this section.

**Fig. 1.**
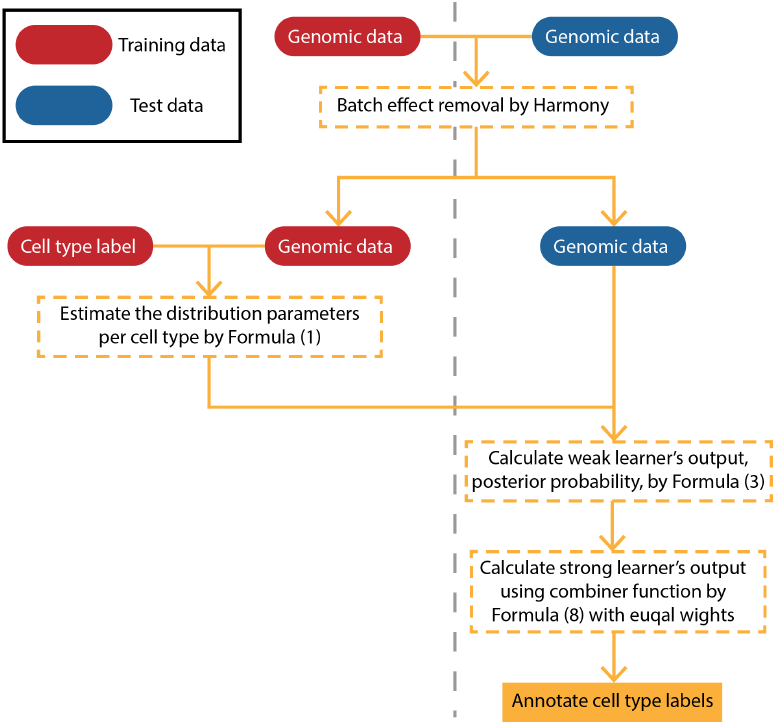
Workflow of scAnnotate on dataset with at most one rare cell population (at most one cell population less than 100 cells). The vertical gray dashed line separates training data (left) and test data (right) information.

**Fig. 2.**
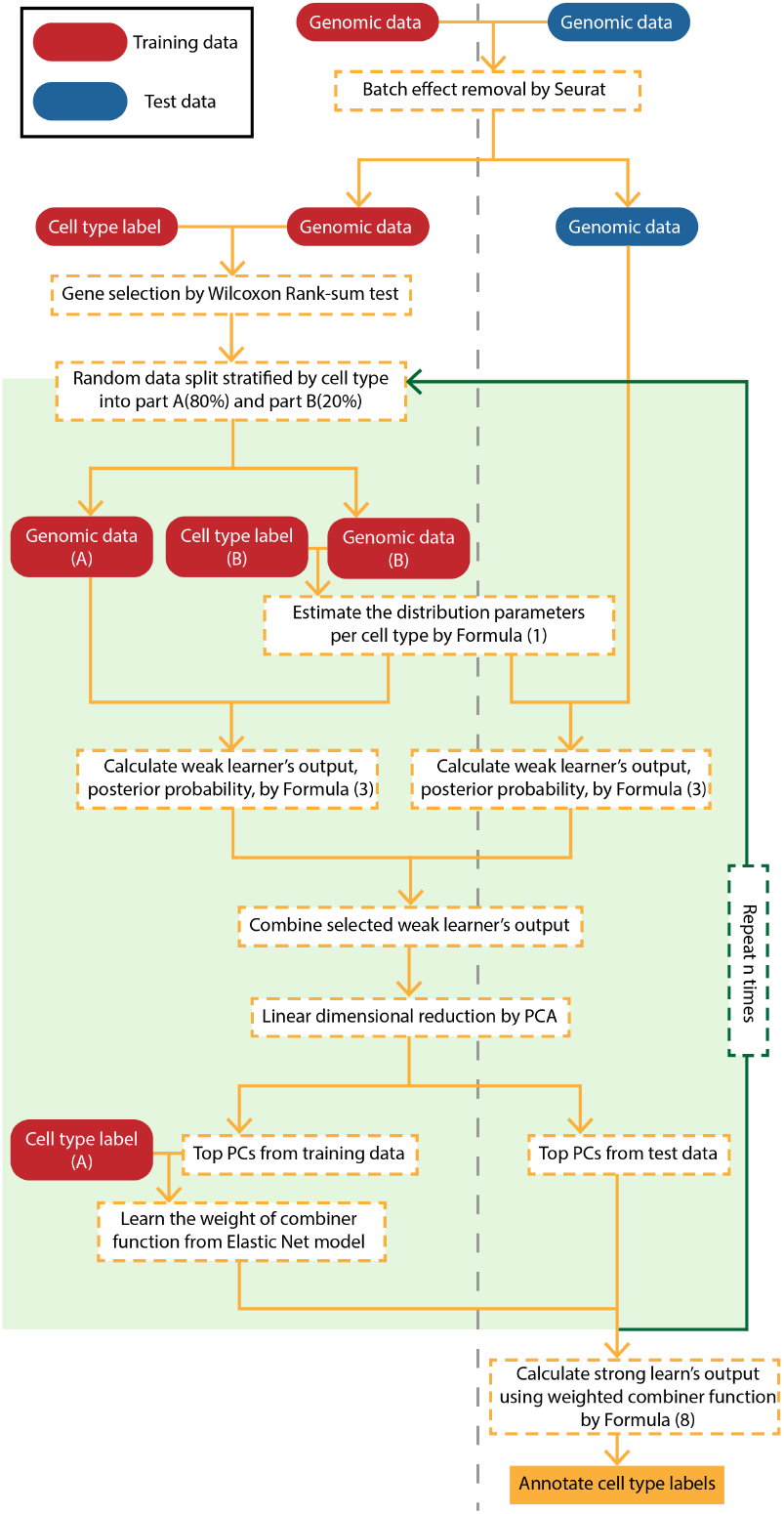
Workflow of scAnnotate on dataset with at least two rare cell populations (at least two cell populations less than 100 cells). The vertical gray dashed line separates training data (left) and test data (right) information.

### 2.1 Batch effect removal

When building a supervised machine learning model for cell-type annotation, batch effects often create differences between the training and testing data. We, therefore, believe that removing batch effects will make the model learned from the training data more suitable for annotating cells in the test data. For example, scPred (Alquicira-Hernandez *et al*., 2019) has batch effect removal as a built-in optional step. Following this idea, we suggest using batch effect removal as a data preprocessing step unless users strongly believe that their training data and test data are similar enough to each other. scAnnotate removes batch effects using the Seurat package (Hao *et al*., 2021) for data with at most one rare cell population or the Harmony package (Korsunsky *et al*., 2018) for data with at least two rare cell populations. Both packages are recommended by Tran *et al*. (Tran *et al*., 2020) due to their consistently high-quality performances and comparatively low runtimes. The output from the batch effect removal step is used as input for the classification model discussed next.

### 2.2 Mixture model for the expression level of a given gene in a fixed cell type

For the *j*th (selected) gene in the type-*i* cell, we propose a mixture model *F*_*ij*_ for its expression level

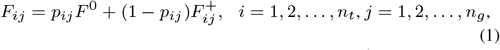

where *F* ^0^ is the degenerated distribution at 0, 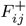 is a distribution supported on (0, ∞), and *n*_*t*_ and *n*_*g*_ are, respectively, the total number of cell types and the total number of genes selected for use in classification. The *p*_*ij*_ and (1 − *p*_*ij*_) are the mixing proportions for *F* ^0^ and 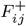, respectively. Model (1) includes commonly used zero-inflated models such as the zero-inflated Poisson model as special cases, but it offers more flexibility than such zero-inflated models as it models the proportion of zeros and the distribution of the positive expression levels 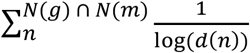 separately. In particular, under (1), all distributions supported on (0, ∞) or a subset of (0, ∞) may be used to model 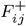. In situations where no good parametric models for 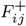 are available, we may also specify 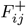 nonparametrically.

### 2.3 Mixture model for the expression level of a given gene, its estimation and prior specification

Let π_*i*_ be the prior probability that a randomly selected cell is of type-*i* where 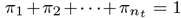. The (prior) distribution of the expression level of the *j*th gene of this cell is the following mixture distribution

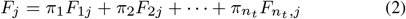

where the *F*_*ij*_ are defined in (1). We estimate *F*_*ij*_ and *F*_*j*_ using training data as follows. Suppose the training data contains *n*_*i*_ independent type-*i* cells. Then, there are *n*_*i*_ independent observations for the *j*th gene in type-*i* cells. Let 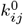 be the number of zeros among these *n*_*i*_ observations and 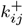 be the number of positive observations so that 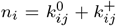 Then, we may estimate the mixing proportions *p*_*ij*_ in (1) with

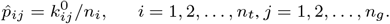

To estimate parameters of the distribution 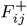, we use the 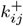 positive observations. For example, if 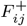 is assumed to be a lognormal distribution, then we can find the maximum likelihood estimates for its parameters using the 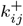 positive observations.

The prior probabilities π_*i*_ depend on the application at hand. In the absence of information for determining these probabilities, we recommend the uniform prior π_*i*_ = 1/*n*_*t*_. We have also used the observed proportions 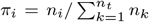 which is reasonable when the training sample is a random sample from the population of cells of all types.

### 2.4 Weak learner based on the mixture model of a single gene

To classify a future cell of unknown type into one of the *n*_*t*_ types with its *n*_*g*_ gene expression data, we first use the *n*_*g*_ genes one at a time to perform the classification. This leads to *n*_*g*_ weak learners.

Let *X*_*j*_ be the expression level of the *j*th gene of the cell. Then, since the type of the cell is unknown, *X*_*j*_ ∼ *F*_*j*_ in (2). Let *x*_*j*_ be the observed value of *X*_*j*_. The posterior probability that the cell is of type-*i* is

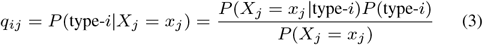

for *i* = 1, 2, …, *n*_*t*_, which can be computed by using the estimated *F*_*ij*_ and *F*_*j*_. Specifically, for *x*_*j*_ = 0, we have

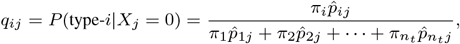

which is the probability that the observed zero comes from a type-*i* cell. For *x*_*j*_ > 0, we have

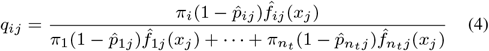

where 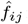 is the estimated probability mass/density function of 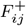. When 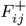 is a continuous distribution, 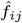 is a continuous density function, so *q*_*ij*_ in (4) is not a real probability but we still call it a posterior probability here for the purpose of classifying the cell. If we only use the expression level of the *j*th gene, we would assign the cell to type-*i*^∗^ where

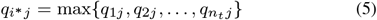

by the rule of the maximum posterior probability. This is a weak learner in the sense that it is based on the expression level of only one gene.

### 2.5 Combiner functions and the strong learner, scAnnotate

With *n*_*g*_ genes, we need to combine the information in the resulting *n*_*g*_ weak learners to obtain an overall classification of the cell based on all *n*_*g*_ genes. To this end, we define an annotation score by combiner function

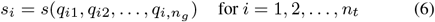

to combine the posterior probabilities for type-*i* from all *n*_*g*_ genes and classify the cell as type-*i*^∗^ where

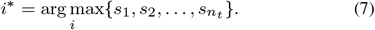

We call steps (1)-(7) mixture model based supervised classification of a cell. For convenience, we will refer to this method as scAnnotate.

#### 2.5.1 Example combiners

We now give several examples of the combiner function (6). The first example is the voting score combiner function

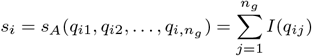

where *I*(*q*_*ij*_) = 1 if 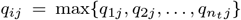 and *I*(*q*_*ij*_) = 0 otherwise. This combiner function essentially counts the number of genes *s*_*i*_ that give the cell a type-*i* classification by the rule of maximum posterior probability shown in (5). With this combiner function, by (5) and (7), scAnnotate classifies a cell as a type-*i* cell if it is most frequently classified/voted as a type-*i* cell by the *n*_*g*_ weak learners.

Another example is the weighted average of the posterior probabilities for type-*i*

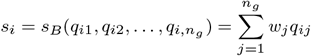

where the weight *w*_*j*_ ≥ 0 and it represents the importance of the *j*th gene. Such importance may, for example, be a quantitative measure of how accurate the classification is when only 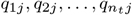 are used to classify cells in the test data through (5). We may also apply the weights to define a weighted version of the voting score combiner,

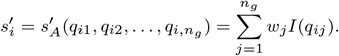

The last combiner example we include here is the weighted sum of log-transformed *q*_*ij*_ -scores defined as

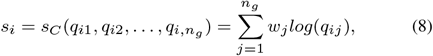

which is equivalent to the product 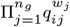. When we use uniform prior π_*i*_ = *P* (type-*i*) = 1/*n*_*t*_ and equal weights *w*_*j*_ = 1 with combiner *s*_*C*_, it is equivalent to 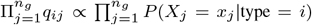, and our ensemble learning reduces to the well-known Naive Bayes Classifier (Rish *et al*., 2001). To see this, the Naive Bayes Classifier uses posterior probability,

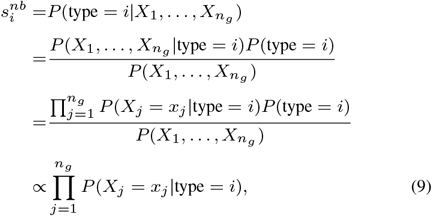

which is equivalent to using 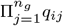 with uniform prior and equal weights. We investigated the combiners given above using multiple real scRNA-seq datasets and empirically found the combiner *s*_*C*_ works best.

#### 2.5.2 Training the combiner

The combiners involve the weights *w*_*j*_, which need to be decided. We assign their values using two different approaches according to the sample size of the training data.

When the training data has at most one rare cell population, we randomly split the training data into two parts. The weights *w*_*j*_ are learned via the following five steps. (1) We use 20% of the cells to estimate the parameters of the *F*_*ij*_. (2) We use the estimated distributions to calculate *q*_*ij*_ scores of the remaining 80% of cells. (3) Using the Wilcoxon Rank-Sum test, we filter out genes whose *q*_*ij*_ scores are not highly associated with cell type labels, i.e. retain the top genes with the smallest *p*-values. (4) Using these 80% cells’ *q*_*ij*_ scores as predictors and their corresponding cell types as outcomes, we train an Elastic Net model (Zou and Hastie, 2005) to learn the weights *w*_*j*_. Note, to reduce the number of predictors, we apply PCA to the scores and use PC scores to replace the *q*_*ij*_ scores. Since the Elastic Net model’s result is a linear combination of PC scores, and the PC scores are linear combinations of *q*_*ij*_ scores, the final results are linear combinations of *q*_*ij*_ scores. (5) To avoid sampling bias introduced by random data splitting, we repeat steps (1)-(4) for 10 times with different random splits, and use average weights learned from the 10 models as the final weights of combiner *s*_*C*_.

When the training data has at least two rare cell populations, we do not have enough data to estimate parameters of all *F*_*ij*_ and at the same time train a model to learn the weights *w*_*j*_ for the combiner. Specifically, a rare cell population has less than 100 cells, as defined for this study. Since we only use 20% of training data to learn the distribution parameters, if we also have to model the weights *w*_*j*_ for the combiner, we cannot sufficiently estimate the parameter *F*_*ij*_ for a rare cell population with less than 20 cells. We can make a reasonable prediction when there is only one rare cell population. The well-estimated distribution of other cell populations can draw an excellent boundary to distinguish the only rare cell population from them. However, we cannot distinguish these rare cells from each other when there are at least two rare cells. In this case, we use all training cells to estimate the parameters of *F*_*ij*_, and assume equal weights *w*_*j*_ = 1 for all *j*.

### 2.6 Implementation of scAnnotate

Classification using scAnnotate depends on three key components: [I] the model for 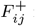 in (1), [II] the prior probabilities π_*i*_ in (2), and [III] the combiner function in (6). For [I], we may use for example Negative Binomial distribution, Exponential distribution, lognormal distribution, or in situations where no good parametric models for positive expression levels are available, a nonparametric measure (see Section 2.7). Due to the usually huge number of combinations of *i* (cell types) and *j* (genes), it is not practical to model each individual 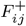 separately, so we assume that distributions of all 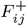 are of the same type, for example, all lognormal; and they can only differ in their parameter values. For [II], we use either a uniform prior or the observed proportions as discussed in Section 2.3. For [III], we may use one of the three combiner functions given above.

In real applications, we recommend using several combinations of the three components to build several classifiers with the training data, and then use test data to evaluate the performance of these classifiers using their *F*_1_ scores to identify the optimal combination with the highest *F*_1_ score. For real data, correct specifications of the components are unknown, so optimizing the combination by trying several combinations and choosing the best one protects scAnnotate from serious misspecifications of its components. For examples that we have tried, we found that the combination of the lognormal distribution, uniform prior and combiner *s*_*C*_ often has the best or second best performance. For simplicity of presentation, this combination is used in all examples in the next section.

Note that in cross-species and cross-platform studies, we need to apply batch effect removal techniques to preprocess the datasets and the processed datasets contain no zeros. For such processed datasets, the mixture model *F*_*ij*_ in (1) for the *j*th gene of a type-*i* cell reduces to 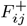 as the proportion of zero *p*_*ij*_ = 0. The mixture model *F*_*j*_ in (2) remains unchanged. The implementation of scAnnotate also remains the same.

### 2.7 Nonparametric depth measure

The true distribution of gene expression can be very complex. We may be unable to find, from the set of commonly used parametric distributions, a suitable one for modelling 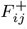. The assumption that distributions for 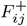 of all genes are the same kind and that they can only differ in parameter values may also be too strong. To deal with these issues, when the sample size is large, we suggest using a nonparametric depth measure for 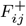 which is totally free of any parametric assumptions. Specifically, we may replace the estimated density function 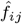 with a depth measure when computing the posterior probability in (4). We now illustrate this point with the use of the halfspace depth measure (Liu *et al*., 1999).

For a fixed gene of a fixed cell type, suppose there are *m* non-zero expression data points from the training data set *x*_1_, *x*_2_, …, *x*_*m*_. Let *x*^∗^ be the expression level of that gene of the cell to be classified. The halfspace depth of *x*^∗^ measures how consistent *x*^∗^ is with the sample *x*_1_, *x*_2_, …, *x*_*m*_. It is defined as follows. First rank the *m*+1 observations in the augmented sample *x*_1_, *x*_2_, …, *x*_*m*_, *x*^∗^ and denote by *r*(*x*^∗^) the rank of *x*^∗^. Then, the halfspace depth of *x*^∗^, *h*(*x*^∗^), is given by

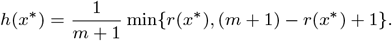

To see how *h*(*x*^∗^) measures the consistency of *x*^*∗*^ with the data, when *x*^∗^ is the smallest or the largest of the augmented sample, *h*(*x*^∗^) has its minimum value of 1/(*m* + 1), so a low *h*(*x*^∗^) value indicates *x*^∗^ is not consistent with the data in that it is an extreme value. On the other hand, when *x*^∗^ is the median of the augmented sample, *r*(*x*^∗^) = (*m* +1)/2 and *h*(*x*^∗^) reaches its maximum value of 1/2, so a large *h*(*x*^∗^) value (close to 1/2) indicate *x*^∗^ is consistent with the data. As such, it may be used to substitute 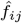 in scAnnotate. Using this depth measure protects scAnnotate from severe misspecification of the model for 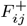. In simulation studies, scAnnotate based on the halfspace depth outperforms scAnnotate with severely misspecified 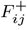 in terms of accuracy. On the other hand, the depth measure requires the number of non-zero observations at all genes to be large (otherwise, the depth measure is too discrete to be a useful replacement for 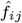) and is more computationally intensive.

To evaluate the performance of scAnnotate, we conduct a benchmark study to compare it against nine other scRNA-seq annotation methods based on supervised machine learning algorithms, including scID (Boufea *et al*., 2020), scClassify (Lin *et al*., 2020), SingleCellNet (Tan and Cahan, 2019), scPred (Alquicira-Hernandez *et al*., 2019), CaSTLe (Lieberman *et al*., 2018), SingleR (Aran *et al*., 2019), CHETAH (de Kanter *et al*., 2019), scmapCluster and scmapCell (Kiselev *et al*., 2018). The parameters of these methods are chosen according to the suggestions in their vignettes or the software default settings; we note that all of the benchmarked methods have a fully automated data-driven approach without requiring previous biological knowledge. Details of the nine methods are listed in Table 1. In our experiments, all models are learned on training data and then applied to annotate cells in the test data. Prior to performing classification, we remove all cells whose cell types do not appear in both the training and test data. To evaluate the classification performance of the benchmarked methods, we compare the predicted labels of the test data with the corresponding true labels. Following the evaluation rule of other annotation method papers (Lieberman *et al*., 2018; Alquicira-Hernandez *et al*., 2019; Lin *et al*., 2020; Zhao *et al*., 2020), we use classification accuracy as our performance criteria. Accuracy in this study is defined as the percentage of correctly annotated cells.

**Table 1.**
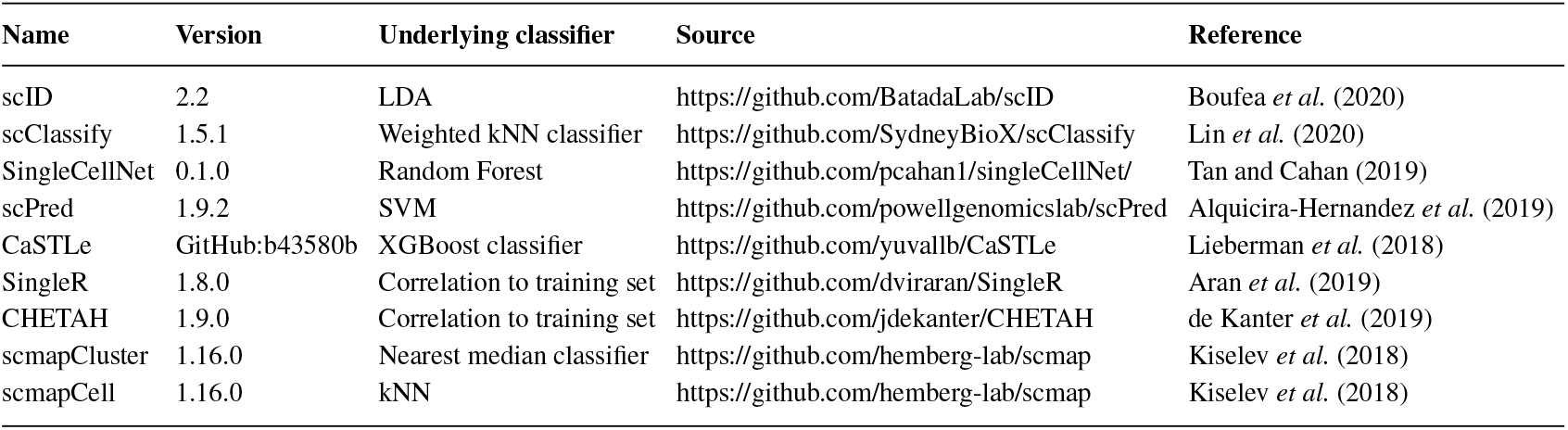
Overview of scRNA-seq annotation methods compared with our method in this evaluation study

We conduct our benchmark study under three situations according to the relationship between training and test data: (1) Training and test data are from the same platform (i.e. obtained from the same sequencing method) and the same species; (2) Training and test data are from different platforms; (3) Training and test data are from different species, human versus mouse.

### 2.8 Datasets and preprocessing

Table 2 summarizes the fourteen publicly available scRNA-seq datasets used in our benchmark study. These data have been used to illustrate the annotation performances of scAnnotate and its nine competitor methods in this section.

**Table 2.**
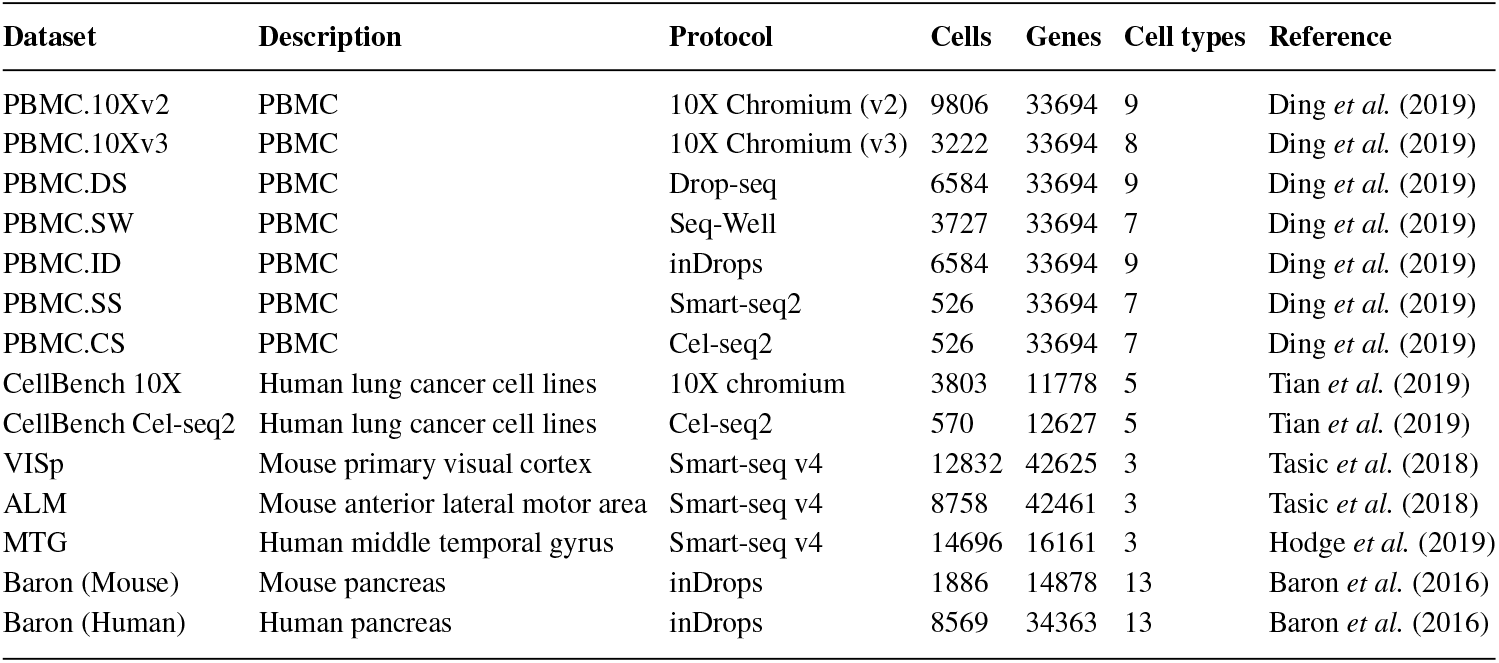
Overview of the datasets used in this study

The human Peripheral Blood Mononuclear Cells (PBMC) scRNA-seq data collection was downloaded from the SeuratData package (Hao *et al*., 2021) with dataset name “pbmcsca” (Ding *et al*., 2019) and consists of seven datasets that were sequenced using seven different methods: 10x Chromium (v2), 10x Chromium (v3), Drop-seq, Seq-Well, inDrops, Smart-seq2, and Cel-seq2. For PBMC cell annotation, we removed all cells labelled as “Unassigned”. Each dataset was then used as training data and all other datasets as test data. This gave us 7 *∗* 6 = 42 distinct pairs of cross-platform datasets. The human lung cancer cell lines data were downloaded from the Zenodo page provided by Abdelaal *et al*. Abdelaal *et al*. (2019). The CellBench 10X dataset was obtained from GSM3618014, and the CellBench Cel-Seq2 dataset was obtained from GSEM3618022, GSM3618023, and GSM3618024. We used both the CellBench 10X dataset and the CellBench Cel-seq2 dataset once as training data and once as test data. This gave us two more distinct pairs of cross-platform datasets.

The mouse and human pancreatic scRNA-seq data were downloaded from the National Center for Biotechnology Information (NCBI) Gene Expression Omnibus (GEO) for GSE84133 (Baron *et al*., 2016). In preparation for cross-species cell annotation, we converted the human gene symbols to mouse ortholog gene symbols using the ortholog table provided by SingleCellNet (Tan and Cahan, 2019). The mouse and human brain datasets were downloaded from the Zenodo page by Abdelaal *et al*. Abdelaal *et al*. (2019). The analysis was limited to common genes between the training and test data. We first used the mouse data as training data and the human data as test data. We then switched the training-testing order and used the human data as training data in order to classify the mouse data. This gave us 1 ∗ 2 + 2 ∗ 2 = 6 distinct pairs of cross-species datasets.

The Baron *et al*. Baron *et al*. (2016) human dataset (GSE84133) was also used for intra-dataset evaluation. We applied stratified sampling (by cell type) to select 80% of the dataset as training data and set the remaining 20% of the dataset as test data. To eliminate the variability in performance evaluation caused by sampling bias, we repeated this experiment ten times with different random data splitting.

The Seurat (version 4.0.5) (Hao *et al*., 2021) package was used for normalization on all raw count matrices. The datasets were normalized using the NormalizeData function with the “LogNormalize” method and a scale factor of 10,000. Since scID was not compatible with log-transformed data, it used the non-transformed normalized data as input. All other methods used the log-transformed normalized gene expression matrix as input.

## 3 Results

### 3.1 Annotation performance evaluation when training and test data are generated from the same species using the same platform

When we randomly split the Baron *et al*. Baron *et al*. (2016) human pancreatic dataset (GSE84133) into training and test data as described in Table 2, scAnnotate (assuming 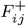 follows a log-normal distribution) classified fourteen pancreatic cell types with a high overall prediction accuracy range of 96.85% to 98.08%. The median prediction accuracy over the ten rounds of classification was 97.69%, and the mean prediction accuracy was 97.69%. Specifically, scAnnotate classified acinar, activated stellate, alpha, beta, delta, ductal, endothelial, epsilon, gamma, macrophage, mast, quiescent stellate, Schwann, and T cells with a mean accuracy of 95.88%, 94.54%, 98.54%, 98.12%, 95.67%, 98.84%, 97.80%, 89.17%, 97.84%, 100%, 100%, 95.10%, 93.33% and 95.00%, respectively. We note that the lower accuracy scores resulted from classifying epsilon and Schwann cells. These two cell types were rare in this dataset; out of the total 8,569 observed cells, epsilon and Schwann cells had only 18 and 13 respective observations. Figure 3 shows the classification accuracy of scAnnotate and competitor methods, including discriminative models scClassify, SingleCellNet, scPred, CaSTLe, SingleR, scmapCluster, scmapCell, CHETAH, and generative model scID. Multiple methods had high classification accuracies when training and test data were similar enough. Among these top performers, scAnnotate ranked third based on prediction accuracy.

**Fig. 3.**
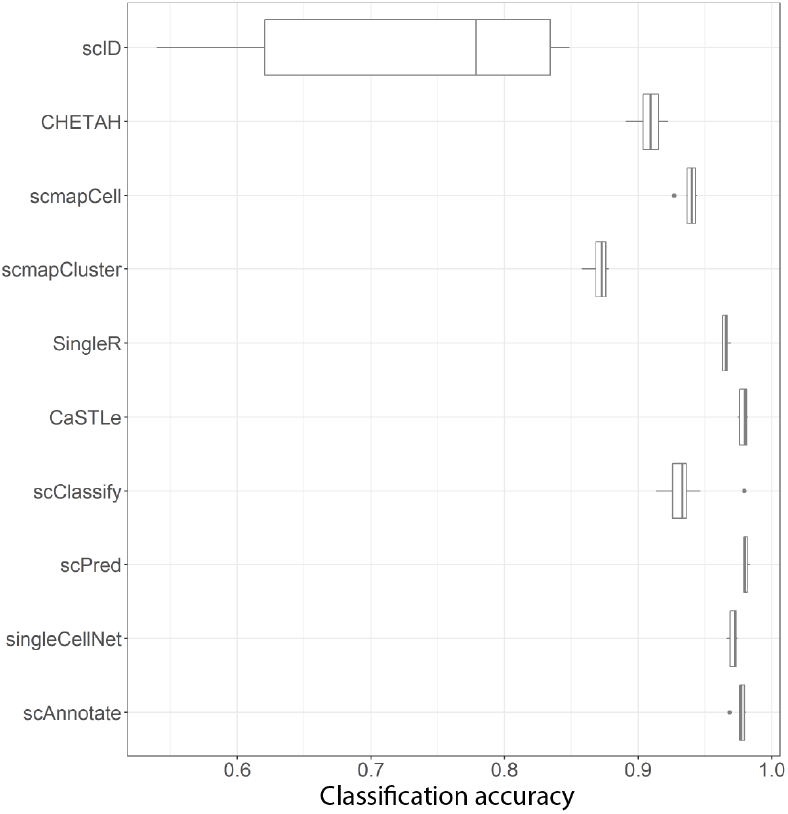
Within-study classification performance of scAnnotate on the Baron et al. Baron et al. (2016) human pancreatic scRNA-seq dataset (GSE84133). The boxplots show the classification accuracies of scAnnotate and nine competitor methods in ten experiments on different random splits of the original data (80% training and 20% testing).

### 3.2 Annotation performance evaluation using cross-platform training data

We used the PBMC datasets (Ding *et al*., 2019; Hao *et al*., 2021) provided by the SeuratData Package and the lung cancer cell lines dataset (Tian *et al*., 2019) to evaluate the ten annotation methods’ classification performances when training and test data are obtained from different scRNA-seq generating platforms. The PBMC data consists of scRNA-seq data generated using 7 different platforms as listed in Table 2, which leads to 42 pairs of cross-platform training and test data. The lung cancer cell lines datasets, 10X and Cel-seq2, provided 2 pairs of cross-platform training and test data. We applied scAnnotate and competitor methods to each of these 44 data pairs and calculated their prediction accuracies.

scAnnotate (assuming 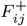 follows a log-normal distribution) had the highest mean accuracy (80.27%) and the highest median accuracy (81.92%), and an overall prediction accuracy range of 39.45% to 100.00%. scAnnotate, singleCellNet and scPred had the highest median accuracies, indicating that they are the top 3 best methods in the overall comparison of the 44 settings. When we look into individual settings, scAnnotate is among the top-ranked most accurate methods in most of the 44 cross-platform settings (Figure 4). When scAnnotate is not the best method, its accuracy is often not much lower than that of the winners. We note that there is a significant drop in performance for scAnnotate when trained on the PBMC 10x-v3 dataset and tested on the PBMC Seq-Well dataset. As shown in the top boxplot of Figure 4, all methods have a significant drop in performance for this cross-platform dataset combination; thus in general, training on the 10x-v3 dataset may produce poor predictions on the Seq-Well dataset. Comparing the performance of each classifier across the different protocols on the lung cancer cell lines, we observe an almost perfect performance for all classifiers.

**Fig. 4.**
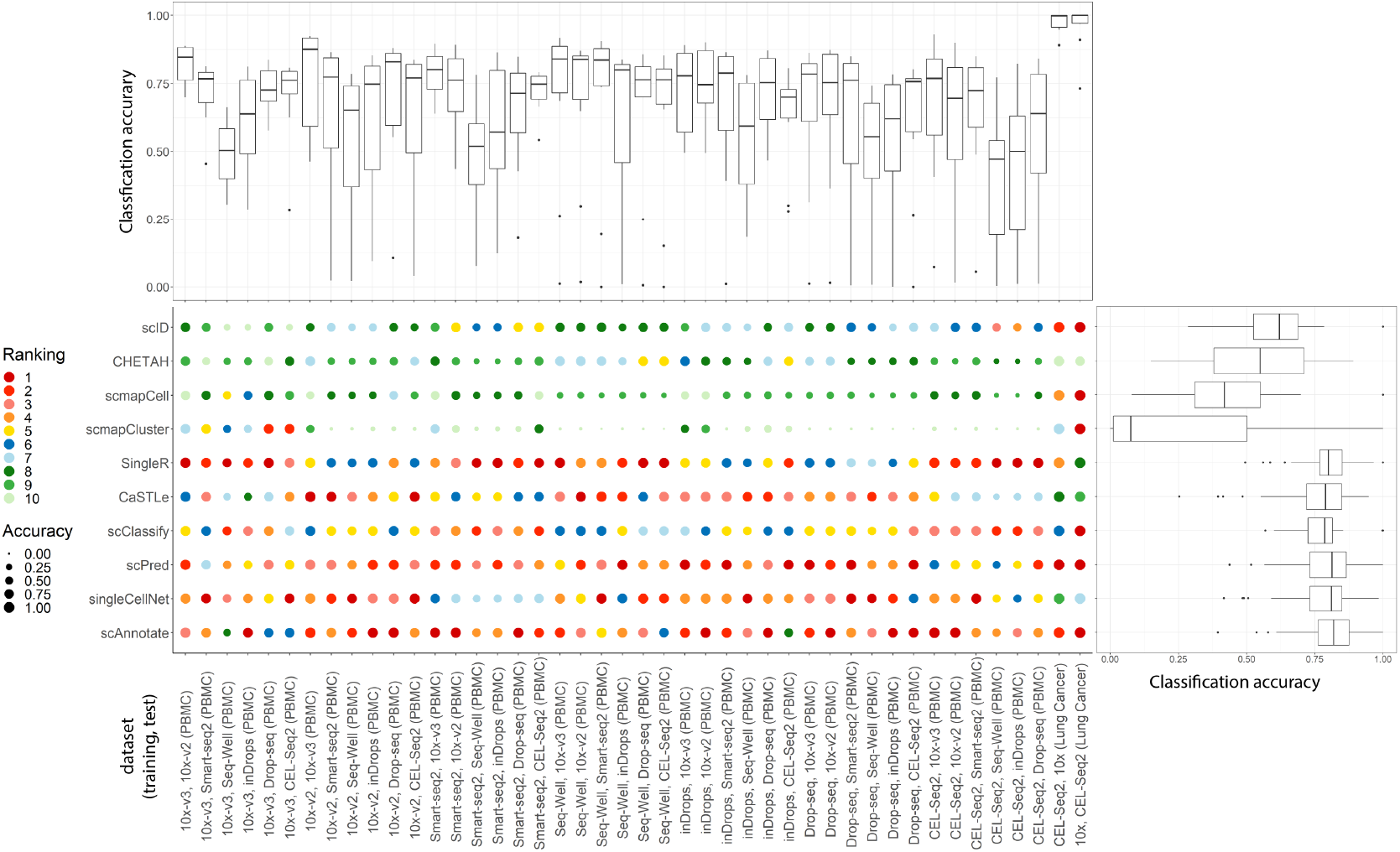
Classification performance of scAnnotate on 44 combinations of cross-platform datasets, as provided by Ding et al. Ding et al. (2019) and Tian et al. Tian et al. (2019). The dot plot shows the comparison results of each individual setting. Each column represents one setting of a training and test data combination. Each circle represents the performance of one method. The colours of circles represent methods’ ranks, and the sizes of circles represent their corresponding accuracies. The boxplot on the right side of the dot plot summarizes the overall comparison of methods under all settings. Each boxplot is constructed using the accuracy values of the given method for the 44 settings. The boxplot on top of the dot plot summarizes the overall performance of classifiers for each experiment. Each boxplot is constructed using the accuracy values of the 10 methods.

### 3.3 Annotation performance evaluation using cross-species training data

We first trained scAnnotate (assuming that 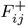 follows a log-normal distribution) and nine other cell annotation methods on the Baron *et al*. Baron *et al*. (2016) mouse pancreatic dataset and predicted ten pancreatic cell types in the Baron *et al*. Baron *et al*. (2016) human pancreatic datasets. scAnnotate ranked first with a prediction accuracy of 92.23%. We then switched the training-testing order and used the human data as training data in order to classify the mouse data. scAnnotate’s rank is third with a prediction accuracy 90.15%.

We then repeated this training/testing process with scAnnotate and the nine competitor methods on the mouse and human brain datasets provided by Tasic *et al*. Tasic *et al*. (2018) and Hodge *et al*. Hodge *et al*. (2019), respectively. Since we utilized two mouse brain datasets and one human brain dataset, switching the training-testing order gave us 4 distinct pairs of datasets. As shown in Figure 5, scAnnotate consistently ranks in the top five best-performing methods for mouse and human brain cell annotation with a mean accuracy 99.15% and median accuracy 99.79%.

**Fig. 5.**
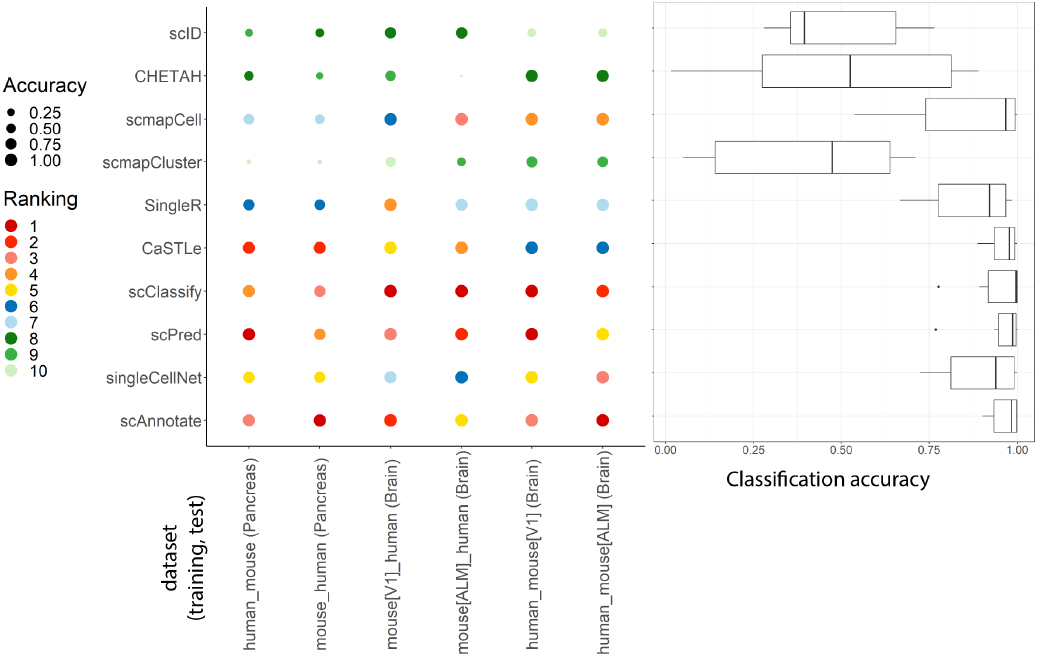
Cross-species classification performance of scAnnotate and nine other methods on six combinations of mouse and human scRNA-seq datasets, provided by Baron et al. Baron et al. (2016), Tasic et al. Tasic et al. (2018) and Hodge et al. Hodge et al. (2019). The dot plot shows the comparison results of each individual setting. Each column represents one combination of training and test data. Each circle represents the performance of one method. The colours of circles represent methods’ ranks, and the sizes of circles represent their corresponding accuracies. The boxplot on the right summarizes the overall comparison of methods under all settings.

### 3.4 scAnnotate can complement other annotation methods

The scAnnotate uses a very distinct modelling approach compared with its competitors, which is the only annotation method that explicitly models the dropout (a critical characteristic of scRNA-seq), and is one of the only two generative classifiers while all others are discriminative classifiers. Therefore, we expect scAnnotate’s annotation of individual cells can be quite different from its competitors, although scAnnotate has similar cohort-level accuracy to other top-performing methods. That is, scAnnotate’s incorrectly annotated cells could be very distinct from its competitors’ incorrectly annotated cells. Being able to complement competitors enables scAnnotate to be used together with other annotation methods, which may lead to improved results. We will further discuss this in the Discussion section.

Next, we look into the detailed annotation results of one data analysis conducted above, as an example, to demonstrate the difference between annotation results of scAnnotate’s generative approach and its competitors (i.e. discriminative methods). We focus on the cross-platform analysis training model on PBMC.SW dataset and tested it on the PBMC.10Xv3 dataset (see Table 2 for dataset details). Of the 2930 cells included in the PBMC.10Xv3 dataset, 2000 cells were correctly annotated by all methods, 44 cells were incorrectly annotated by all methods, and the rest 886 cells were inconsistently annotated by these methods. The mosaic plot of Figure 6 shows the comparison of annotation results of six top-performing annotation methods. Each row represents an annotation method, and each column represents a cell. The grids filled with black color indicate those cells are incorrectly annotated. To highlight the pattern of difference between methods’ annotation results, we only show the results of the top six best-performing methods, and exclude cells that are consistently annotated by all methods. We grouped these cells into 20 clusters using the K-Means algorithm, and re-ordered the cells according to their cluster membership to highlight the pattern. From Figure 6, we can observe an obvious pattern that scAnnotate’s results complement all other methods. In this analysis, SingleR had the best performance with an accuracy of 91.74%; scAnnotate ranked second with an accuracy of 90.14%. If we could let methods correct each other’s errors (e.g. by manually investigating inconsistently annotated cells or building an ensemble model to borrow information among different methods), the ideal accuracy could be improved to 98.50% in this example.

**Fig. 6.**
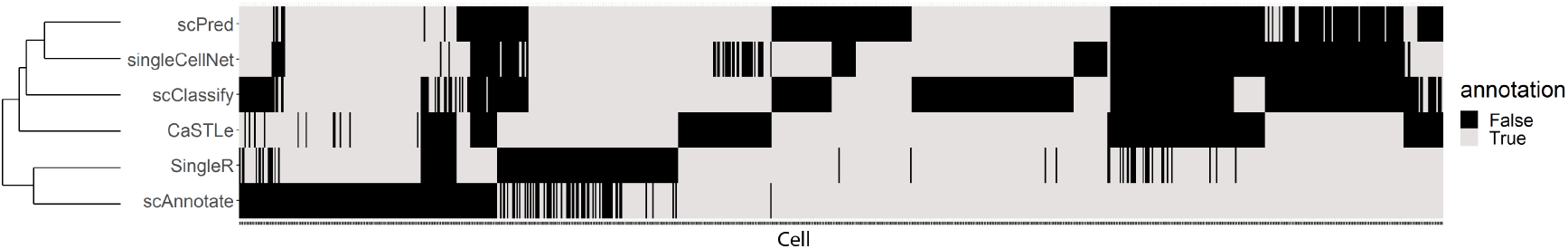
The mosaic plot shows the cells of the PBMC.10Xv3 dataset that are incorrectly annotated by at least one of the top six benchmarked methods (when trained on the PBMC.SW dataset) and correctly annotated by others (scAnnotate, singleCellNet, scPred, scClassify, CaSTLe, SingleR). The dendrogram on the left shows the hierarchical clustering of cells into types by each of the top six methods.

## 4 Discussion

scAnnotate is an analysis framework that consists of three major components. Users can change the components according to their own needs. First, we use two available softwares as a batch effect removal step: Harmony for dataset with at least two rare cell populations, and Seurat for dataset with at most one rare cell population. Many batch effect removal methods can be used to replace them, such as the methods compared in the benchmark study by Tran *et al*. Tran *et al*. (2020). Second, an Elastic Net Zou and Hastie (2005) model is used to learn a combiner function. Users can choose any supervised machine learning model to replace it. For example, when the sample size is large, users can train the combiner function using XGBoost (Chen and Guestrin, 2016), which is faster and much more precise than Elastic Net in this situation. Third, the distributions used in weak learners can also be changed. We will discuss this next.

After investigating many candidate distributions used in weak learners, we recommend using the lognormal distribution as the non-dropout component of our mixture model. Besides lognormal, we added two alternative distributions in our software for particular usages. The first alternative distribution is the exponential distribution. This distribution has only one parameter, and hence is most suitable for rare cell type detection with small and unbalanced sample sizes. The second alternative is the non-parametric distribution estimated using a depth function approach, used for extra-large sample size problems. We believe that all distributions oversimplify the complex truth of gene expression. Hence, a non-parametric approach can better estimate the complex true distribution when the sample size is big enough. We believe that this alternative will be more useful in the future as the sample size of scRNA-seq data continues to grow rapidly over the years.

Differential expression analysis uses genes one at a time; most DE methods, therefore, focus on modelling gene expression distribution to best utilize key features of genomic data. In contrast, annotation analysis needs to use all genes to make decisions. Most cell annotation methods do not model the distribution of genomic data, since it is hard to model the joint distribution of many genes. We address this challenge using an ensemble approach. We build classifiers using each gene separately and ensemble them using a combiner. This approach has three advantages in computing. First, by modelling each gene separately, the number of parameters in our model is linear in the number of genes, which successfully avoids the curse of dimensionality. Second, the estimation of distribution parameters in the training data and the evaluation of posterior probability in the test data often have close-formed formulas, rather than the expensive iterative approximation used by many other types of classifiers. Third, when necessary, such calculations for tens of thousands of genes can be done in parallel to further reduce computing time.

While scAnnotate is developed as a stand-alone tool for cell annotation, it can also be used with other methods to correct each other’s errors. According to the “No free lunch theorem” Wolpert and Macready (1997), there is no single best machine learning algorithm for predictive modeling problems. All methods have their particular advantages and disadvantages. Hence, different annotation methods are expected to make errors on different sets of cells. At the end of the Results section, we demonstrate that scAnnotate’s annotation errors are very different from its competitors’. In practice, users could use scAnnotate and another method to analyze the same data and compare their annotation results. Users could manually investigate the cells annotated inconsistently between scAnnotate and its competitors to improve the accuracy of cell annotation further. When there are enough computing resources and training data, users could also build an ensemble annotator to automatically integrate the annotation results of many methods. In such an ensemble annotator, we believe scAnnotate should play a key role because of its distinct characteristics and ability to complement competitors. Users also could use annotation results of scAnnotate and other methods as input for downstream analyses, and then compare the final results to identify which one makes more sense in biology.

## 5 Conclusion

In conclusion, we introduce scAnnotate, a streamlined process for scRNA-seq data analysis that includes data preprocessing and cell annotation. It is an entirely data-driven automated method that requires no biological knowledge or subjective human decisions (such as pre-specifying the feature genes of cell types). We simultaneously model genes’ dropout proportions and expression levels via a two-component mixture model. We use an ensemble machine learning approach to address the curse of high-dimensionality. We build one weak classifier using each gene, and use a combiner function to integrate all weak classifiers into a single strong classifier. Using multiple real scRNA-seq benchmark datasets, we show that scAnnotate accurately identifies cell types when training and test data are from (1) the same platform and species, (2) different scRNA-seq generating platforms, and (3) different species (specifically, mouse to human). Compared to other supervised machine learning methods, scAnnotate provides top-tier cell classification performance.

## Data and software availability

The data used in this study are all publicly available. The details about how to access these public data are described in the Materials and Methods section. The code to reproduce all of the analyses presented in our study is available on GitHub: https://github.com/ubcxzhang/scAnnotate_reproduce. An open-source implementation of scAnnotate is available as an R package from CRAN.

## Acknowledgements

The authors thank the anonymous reviewers for their valuable suggestions. This research was enabled in part by computing resources support provided by WestGrid (www.westgrid.ca) and Compute Canada (www.computecanada.ca).

